# Predictions of DNA mechanical properties at a genomic scale reveal potentially new functional roles of DNA-flexibility

**DOI:** 10.1101/2023.04.06.535841

**Authors:** Georg Back, Dirk Walther

## Abstract

Mechanical properties of DNA have been implied to influence many its biological functions. Recently, a new high-throughput method, called loop-seq, that allows measuring the intrinsic bendability of DNA fragments, has been developed. Using loop-seq data, we created a deep learning model to explore the biological significance of local DNA flexibility in a range of different species from different kingdoms. Consistently, we observed a characteristic and largely nucleotide-composition-driven change of local flexibility near transcription start sites. No evidence of a generally present region of lowered flexibility upstream of transcription start sites to facilitate transcription factor binding was found. Yet, depending on the actual transcription factor investigated, flanking-sequence-dependent DNA flexibility was identified as a potential factor influencing binding. Compared to randomized genomic sequences, depending on species and taxa, actual genomic sequences were observed both with increased and lowered flexibility. Furthermore, in *Arabidopsis thaliana*, crossing-over and mutation rates, both *de novo* and fixed, were found to be linked to rigid sequence regions. Our study presents a range of significant correlations between characteristic DNA mechanical properties and genomic features, the significance of which with regard to detailed molecular relevance awaits further experimental and theoretical exploration.

## Introduction

The structure and mechanical properties of double-stranded DNA (dsDNA) have been of interest since its discovery in 1869 (1). Since then, besides its primary double-stranded helical structure, also known as B-DNA, a variety of different secondary structures of DNA have been discovered. Different helical forms, such as A-DNA and Z-DNA, DNA G-quadruplex and cruciform-DNA have been shown to have a wide range of biological functions (2–5). In addition to the existence of distinct conformational variants, basic mechanical properties of the dominating B-form of genomic DNA influence every aspect of genomic DNA maintenance within cells, its functions, and interactions with other molecules, mainly proteins. The eukaryotic genome is packaged in the form of nucleosomes, which is formed by a ∼150bp stretch of DNA wrapped around a histone protein complex. Nucleosome formation has been linked to flexibility of genomic DNA, with histones binding to more flexible DNA regions, while linkers and nucleosome-depleted regions (NDR) have been reported to be more rigid (6–8). Specifically, NDRs have been connected to long poly-dA:dT tracts with rigid structure resistant to sharp bending (9–11). Variable DNA flexibility has also been discussed in the context of nanotechnological applications (12).

With regard to interactions of genomic DNA with proteins, transcription factor (TF) binding has long been considered connected to DNA shape. Both detection capabilities of proteins for specific shape properties of DNA as well as bending of the DNA upon protein binding have been reported (13–15), reviewed in (16). A recent publication succeeded in predicting DNA-protein interaction by explicitly considering DNA shape as a determinant feature (17). A puzzling observation with regard to TF-binding to genomic regions has been that some loci with canonical TF-binding motifs present were found to be bound by TFs, while others were not. Recently, it has been suggested that the flanking regions of the binding motif of TFs are key in determining binding efficiency. Since there is more sequence variation in these flanking regions than in the consensus motifs, it has been suggested that mechanical properties, specifically their flexibility, may play a major role in defining the binding site (18–20).

The mechanical flexibility of dsDNA is determined by the constraints of its backbone, high charge density (negatively charged phosphate groups), as well as its sequence of bases and their stacking interactions (21, 22). dsDNA was considered among the most rigid biompolymers with a persistence length of ∼50nm, or approximately 150bp (21). However, recent studies showed that short DNA sequences under 100bp are able to loop spontaneously, indicating that DNA is much more flexible on a short scale than previously assumed (23–25). The underlying sequence determinants has hence been intensively researched, for review see (26).

Based on the potential for looping of short DNA sequences, several assays for measuring DNA flexibility of short sequences have been developed (for a recent review, see (27)). Loop-seq, developed recently by Basu *et al.* (28), uses a sequencing-based approach, allowing a high-throughput measurement of DNA mechanical properties. For this method, a library of DNA fragments flanked by two adapter sequences is immobilized and nicked. Then, the sequences are allowed to loop for a specific amount of time. Afterwards, unlooped DNA is digested, the library is amplified, sequenced, and compared to a control treated identically, but without digestion. Overrepresented sequences have a short looping time, meaning they are more flexible, while underrepresented sequences are more rigid. The natural logarithm of the ratio of the relative population of a sequence in the sample pool to that in the control was termed cyclizability, with a low value representing rigid, and a high value flexible sequences.

The study by Basu et al. confirmed many previously observed effects of DNA mechanics, like being a key factor in nucleosome positioning. In addition, they were able to show that codon usage might be influenced by DNA mechanics. Furthermore, their study also provided a large dataset linking sequence and measured cyclizability, allowing the development of machine learning prediction tools. An accurate computational prediction model may find many applications and may even potentially replace experimental methods, while also leading to a higher base-pair resolution. Accordingly, attempts have been made and high prediction accuracies have been reported (29, 30).

In this study, a deep learning CNN/LSTM model, CycPred, was developed and utilized to further explore the biological functions of local DNA flexibility. Our work builds on the previously published studies on experimentally measured (28) and computationally predicted DNA cyclizability scores (29, 30). We investigated the significance of mechanical flexibility in several species from different kingdoms, including plants, with regard to magnitude relative to random expectation, underlying sequence determinants, sequence range, and genomic features, such as transcription start sites, TF-binding sites, single nucleotide polymorphisms (SNPs) and *de novo* mutations, DNA-methylation-, and crossing-over sites. We show that mechanical properties may indeed play a crucial role in determining the sequence-structure-function relationships of genomic DNA.

## Materials and Methods

### Utilized Datasets

For training and validating the computational prediction models, the datasets for intrinsic cyclizability in yeast as reported by Basu et al. (28) were utilized. All intrinsic cyclizability datasets consisted of sequences of length 100bp, of which 50bp are actual probe-sequence, with 25bp flanking sequences on either side. The utilized sequences were the 82,404 probe-sequence windows with step size 7bp for chromosome V, 19,907 sequences for the nucleosome library, 82,368 sequence windows with step size 7bp for the nucleosome tiling library, and 12,472 sequences for the library containing random sequences.

The *Saccharomyces cerevisiae* genome S288C and its annotation was downloaded from the SGD database https://www.yeastgenome.org/. The genome of *Arabidopsis thaliana*, TAIR10, and its annotation was downloaded from https://www.arabidopsis.org/. The genome and annotation of *Chlamydomonas reinhardtii* was downloaded from Phytozome (https://phytozome-next.jgi.doe.gov/). The genomes and annotations of all other species were downloaded from https://www.ncbi.nlm.nih.gov/.

Methylation calls for *Arabidopsis thaliana* were taken from the 1001 genome project (31), as were the SNP and short indel calls (32). With regard to *de novo* mutations, calls as reported by Monroe et al. (33) were utilized.

Transcription factor binding site (TFBS) motifs were obtained from JASPAR (34), and DAP-seq data, indicating transcription factor binding events to “naked” DNA, for *Arabidopsis thaliana* from O’Malley et al. (35).

The Spo11-oligo-seq Arabidopsis dataset utilized in this study was published by Choi et al. (36) and kindly provided by the study authors. It is reported in the from of log2(spo11-oligonucleotides/gDNA), with spo11-olignucleotides representing number of reads mapped at a genomic position, and gDNA representing number of single-end reads from genomic DNA mapped to the same position as control. The data was mapped to the Col-Cen genome (37), which was obtained from https://github.com/schatzlab/Col-CEN.

### Design and training of the deep learning model

The Python package tensorflow-gpu 2.6 was used to create the deep learning models for predicting the measured cyclizability values (28) based on DNA-sequence (Figure 1). For optimizing the model architecture, the package kerastuner 1.04 was utilized.

**Figure 1.**
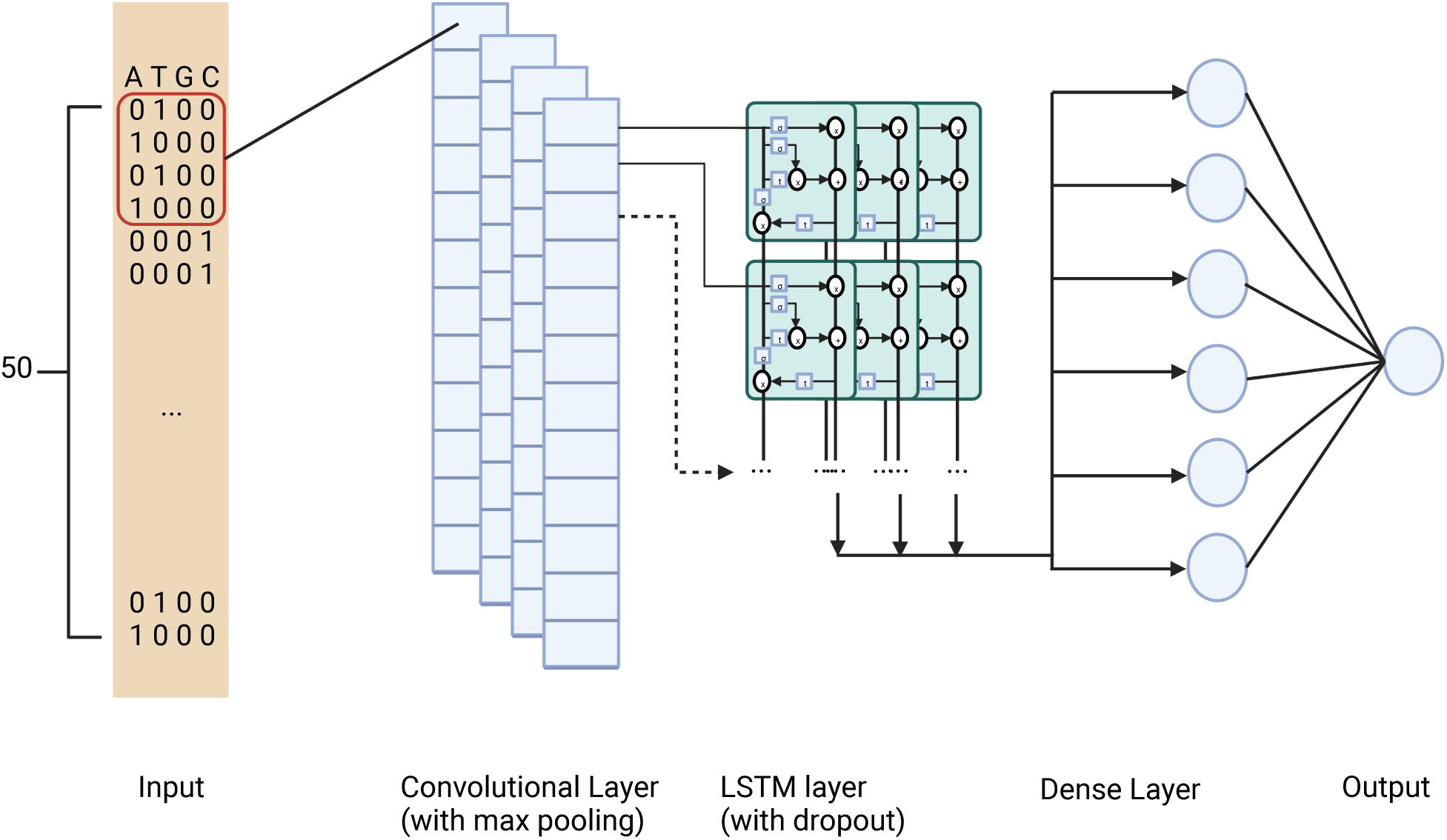
Architecture of the prediction model.

In all intrinsic cyclizability datasets, adapter sequences (25 bases at either terminus) were removed, resulting in sequences of length 50bp. The tiling library was split into training and test sets with a 70% / 30% split. As the dependent variable, the intrinsic cyclizability score denoted as C0 was used.

As the loss function the mean squared error (MSE) was applied. The model utilizes 1D convolution layers with max pooling, followed by an LSTM layer with dropouts. The last hidden layer is a dense layer followed by the output layer (Figure 1). Due to the discrepancy between cyclizability of forward and reverse sequences, which was also reported in the original paper (28), both the forward and reverse-complement of each sequence windows were predicted and the mean was reported as cyclizability value.

For identification of potential motifs identified by the model, we used deeplift (38) for calculation of importance scores, and modisco (39) for the identification of potential motifs based on those importance scores.

### Calculation of Cohen’s d effect size

Cohen’s d effect sizes were calculated as follows:

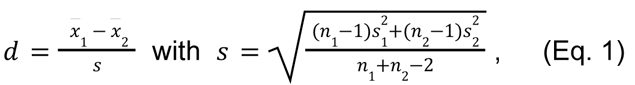

where *x_1/2_* are the respective means of two distributions 1 and 2, and s_1/2_ their associated

standard deviations.

### Analysis of sequences around transcriptions start sites (TSSs)

TSS positions were determined using available genome annotations (gff files), with the position of the first base of the 5’ UTR taken as the TSS. For each gene, only one isoform, isoform 1, if available, was chosen.

Starting from position -525bp (with the negative sign indicating position towards the 5’ of DNA, i.e. “upstream”) from the TSS and ending at position +525bp, the 1050bp long sequence results in 1000 windows of length 50bp. For each window, cyclizability was predicted using the trained model. Each position in the respective profile plots represents the mean of all windows with this position as their 25th base, where the mean is taken across all determined TSSs in the respective genome.

For investigation of the effects of nucleotide/dinucleotide composition on the cyclizability values, we applied a shuffling and a sequence sampling approach. For shuffling, each above-mentioned 50bp sequence window was either randomly shuffled or shuffled with the fasta-dinucleotide-shuffle algorithm (40). In essence, sequences were shuffled “horizontally”). For sampling, the nucleotide-and dinucleotide-probability at each position of the 1050bp long sequences were calculated based on actual sequences around all TSSs. Nucleotide-based sampling involves simply drawing nucleotides according to their probability at each position. For dinucleotide sampling, a random sequence was generated by adding nucleotides to a growing sequence, initialized by a single base. Added nucleotides are drawn based on the conditional probability of the four nucleotides on the preceding base, initialized at the first position according to the observed base probability at this site. Conditional probabilities were determined based on all actual sequences around TSSs. The latter can be referred to “vertical sampling” across all TSS-centered sequences with consideration of conditional probabilities, similar to a first-order hidden Markov model.

### Statistical analysis of trends in cyclizability at a genomic scale

For each organism, we sampled 50bp windows in 50bp steps from the whole genome. Windows containing unresolved bases were removed. Each of the windows was randomly sequence-shuffled five times respectively, and the difference between the prediction of the actual genomic sequence and the predictions of the shuffled sequences was reported. A resulting value below zero indicates that the genomic sequence has a lower predicted cyclizability, and is thus “stiffer” than expected for randomly shuffled versions of the same sequence, a value above zero indicates the opposite, i.e. more flexible than expected.

### Analysis of *de novo* mutations, fixed SNPs, and methylated positions in *Arabidopsis thaliana*

Sequence windows of length 50bp containing methylated positions were compared to randomly shuffled versions (as explained in “Statistical analysis for trends in cyclizability at a genomic scale”).

For both naturally occurring (fixed) SNPs and *de novo* mutations, all windows with a *de novo*/ fixed SNP were sampled, their cyclizability was predicted and compared to the predicted cyclizability of all windows with no reported mutation in the respective dataset.

### Analysis of cyclizability in relation to transcription factor (TF) binding

TF binding was analyzed by comparing bound motifs in the promoter region, defined as 1000bp upstream of the TSS, to unbound occurrences of the same motif. A TF motif was considered “bound” when there was an overlap between DAP-seq region reported for the respective TF and a FIMO (41) search with the motif-PWM (position weight matrix) associated with the respective TF with a q-value < 0.1, and unbound otherwise.

### Analysis of Spo11 oligonucleotides

For each non-zero value of log2(spo11-olignucleotide/gDNA) excluding the centromere region on Arabidopsis chromosome 1, cyclizability for the window centered around the respective position was predicted. For statistical analysis Pearson correlation between Spo11 value and predicted cyclizability was calculated. To compare the top/bottom percentile, both the cyclizability scores as well as the Spo11 values for each position were binned, the positions with overlapping bins counted, and a Fisher exact test was performed.

## Results

We first report on the performance of our computational prediction method with regard to predicting cyclizability, and thus local genomic DNA mechanical flexibility/ rigidity. After having established high accuracy and examining the underlying sequence-determinants of cyclizability, we then apply the model to investigate differences between the genomes of several species from different kingdoms and the potential relevance of local DNA rigidity on a range of different phenomenon, such as transcription initiation and transcription factor binding, and specific genomic sites including methylation sites, variants, and double-strand DNA breaks, and with a particular focus on the genome of the plant *Arabidopsis thaliana*, for which rich annotation and additional genomic-variant information is readily available. Of note, the actual target variable which the model was trained on, is “cyclizability”, i.e. the ability to form closed-loops. As we take this parameter as a surrogate of “flexibility”, we use this term, as well as its logical complement, “rigidity”.

### The CNN/LSTM model shows similar performance as DNAcycP, while being computationally less expensive

Our computational prediction model, termed CycPred, uses 1D Convolutional neural network (CNN) layers with max pooling followed by a LSTM layer and two dense layers (Figure 1).

The model DNAcycP by Li *et al.*(29), published during our work on this project and with identical objective and underlying input datasets, has a similar architecture, utilizing a CNN based Inception-ResNet structure and an LSTM layer.

DNAcycp and our model show a similar performance. In the case of prediction of the 12,742 sequences in the random library, both yielded a correlation between predicted and actual cyclizability score of 0.93 (Pearson correlation coefficient), with a median distance between predicted and actual score of 0.092 for DNAcycP and 0.089 for our model. In case of the chromosome V library data, both models show a correlation of 0.77, with DNAcycP having a median distance of 0.176 and our model of 0.178. A major difference between the two prediction models was the required run time. While a direct performance comparison is difficult, DNAcycP needed between 60x and 80x more processing time for input sequence to our CycPred model. Since many of our analyses require prediction of large sequence sets, all analyses were performed with our prediction model CycPred.

### Pronounced changes of cyclizability around transcription start sites

The study by Basu et al. (28) showed that in yeast, sequence windows located approximately 150bp upstream of the so-called “+1 dyad”, i.e. the position of the first location of high nucleosome occupancy downstream of the TSS, has a pronounced lower cyclizability relative to neighboring sequence locations. Given that this region of significantly lowered nucleosome occupancy is located approximately 20bp-50bp upstream of TSSs, this result was interpreted as an indication of genomic DNA being rigid in locations of low nucleosome occupancy, so-called nucleosome depleted regions (NDRs), thereby facilitating access of transcription factors to their respective target sites in gene promoters, an observation that agreed with previous considerations to this effect (42). By also developing and applying a computational prediction tool, Li *et al.* (29) were able to demonstrate that also in mouse, nucleosome positions in general coincide with increased flexibility. However, unlike in yeast, where a rigid region was observed upstream of the TSS, in mouse, a rigid region was found downstream of the TSS.

To further investigate the potential role of DNA mechanical properties in transcription initiation, we examined the characteristic profile of cyclizability around TSSs in eight different species, three plants (*Arabidopsis thaliana* (Ath)*, Orzya sativa* (Osa)*, Nicotiana tabacum* (Nta)), one single-cell alga (*Chlamydomonas reinhardtii* (Cre)), yeast (*Saccharomyces cerevisia*e (Sce)), two mammals (*Mus musculus* (Mmu)*, Homo sapiens* (Hsa)), and the invertebrate *Caenorhabditis elegans* (Cel) (Figure 2). Noticeably, in all species, pronounced changes of cyclizability near the TSS are apparent. Six of the eight species (all but Sce and Cre) show very similar profiles with a pronounced and TSS-downstream-sided dip of cyclizability. In yeast (Sce), a pronounced dip was found ∼ 0-150bp upstream of the TSS, thus approximately where it had been reported previously (28, 43). In addition to the observed dips, regions of increased cyclizability, noticeably flanking the dips, are also apparent (upstream of the dip: Ath, Osa, Nta, Cel, or symmetrical around it: Sce). The pattern of change of cyclizability in the alga Cre stands out, exhibiting a cyclical behavior.

**Figure 2.**
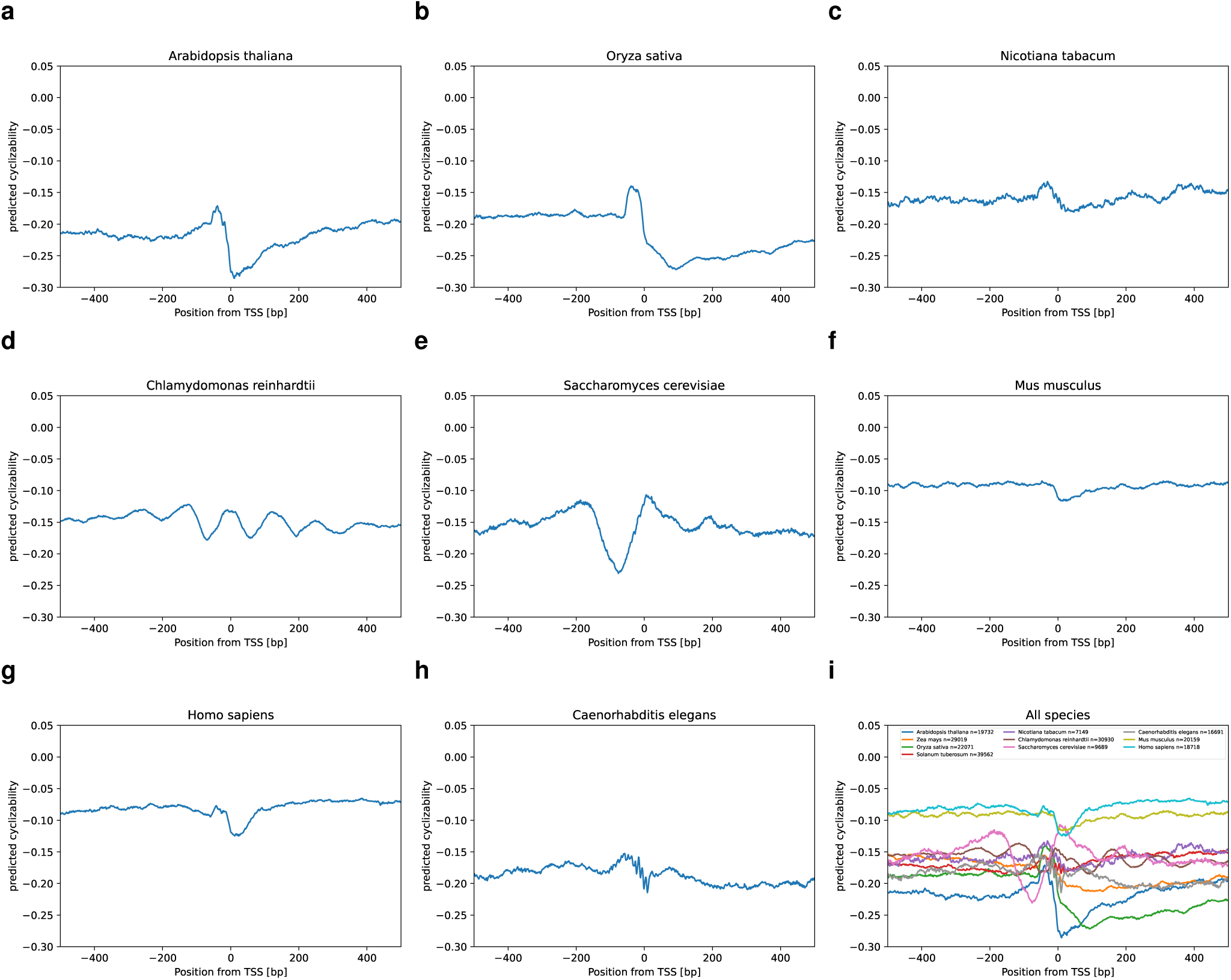
Comparison of predicted cyclizability around transcription start sites (TSSs) of different species. Mean predicted cyclizability around the TSS of all annotated genes containing a 5’ UTR of **(a)** *Arabidopsis thaliana*, **(b)** *Oryza sativa*, **(c)** *Nicotiana tabacum*, **(d)** *Chlamydomonas reinhardtii*, **(e)** *Mus musculus*, **(f)** *Homo sapiens*, **(g)** *Caenorhabditis elegans*. **(h)** Summary of all species.

### Autocorrelation of cyclizability shows no large-scale patterns

As we observed a characteristic and pronounced change of cyclizability locally around TSSs (Figure 2), we tested whether cyclizability exhibits long-distance patterns in the genome. Using the genome of *Arabidopsis thaliana* as a test species, we computed the autocorrelation function of cyclizability for chromosome 1. With autocorrelation, the persistence length of similar cyclizability as well as the presence of periodic patterns would be detectable. However, there was no long-range pattern discernible (Figure 3 a,b). Autocorrelation decreased rapidly with positional distance (lag) from the reference position, dropping off to near zero shortly before 50 bp, incidentally the size of windows used to predict local cyclizability. This shows that in *Arabidopsis thaliana* there is no global (<10,000bp) periodic pattern of local DNA flexibility and that, furthermore, cyclizability is a local property and is generally not maintained at similar levels across larger genomic regions. However, this conclusion does not contradict earlier findings that cyclizability exhibits a periodic behavior in nucleosome (dyad) regions (29), but may indicate that nucleosome positioning is not periodic.

**Figure 3.**
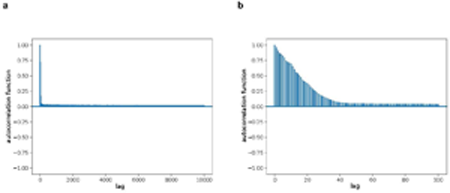
Autocorrelation shows no long-range patterns. Autocorrelation function of predicted cyclizability *Arabidopsis thaliana* chromosome 1, window size 50bp, step-size 1bp, **(a)** with maximum lag of 10000bp, **(b)** with a maximum lag of 100bp.

### Dinucleotide composition is a driving factor in local DNA-rigidity

Previous studies (29, 30) as well as the model created in this study showed high accuracy of computationally predicting cyclizability values of the loop-seq data (r=0.94, Pearson correlation of actual vs. observed cyclizability), suggesting that there is a set of “rules” determining the rigidity of DNA that the model was able to learn and that rigidity is determined by sequence. Previous studies concluded that base composition is not a good indicator of DNA rigidity, except for mono-base or two-base repeats (44). This was confirmed by the two existing computational models (29, 30) as well as by our model CycPred. Randomly shuffling random sequences (each base assumed with equal frequency (25%), shuffling at the level of individual bases) leads to a large variance of predictions. Performing 1000 repeats of shuffling of random sequences 1000 times respectively led to a median standard deviation of 0.329 of predicted cyclizability. However, when shuffling at the level of dinucleotides, performing the same experiment on the same set of random sequences, yielded a significantly different set of standard derivations (p < 0.0001, Cohen’s d= -0.64), with a median of 0.293, and thus significantly smaller than obtained for single-base-level shuffled sequences.

To assess the importance of sequence and composition for cyclizability with regard to single bases and dinucleotides, we compared predicted cyclizability values of actual sequence windows to randomized versions, randomizing either at the level of single bases or nucleotides. We employed two different randomization strategies, one dubbed “vertical”, and one “horizontal” shuffling, see Methods.

Applying the “vertical” shuffling protocol, in which random sequences were generated by statistically sampling from the average composition at specific position near TSSs, either on a single or dinucleotide basis, the signal of a characteristic change of cyclizability around TSSs was completely lost (i.e. flat) for single-base-sampling in all three inspected species (*Arabidopsis thaliana*, yeast, and human) (Figure 4a,b,c). By contrast, dinucleotide sampling yielded barely distinguishable cyclizability change profiles compared to the actual profiles (Figure 4 b,d,f). However, with regard to absolute level, species differed. While single-base shuffling led to increased values in *Arabidopsis thaliana* and yeast (Figure 4 a,c), lowered values were obtained in human (Figure 4e). Applying dinucleotide statistical sampling, actual and randomized sequences tracked one another for *Arabidopsis thaliana* and yeast (Figure 4 b,d), but now were increased for human (Figure 4f).

**Figure 4.**
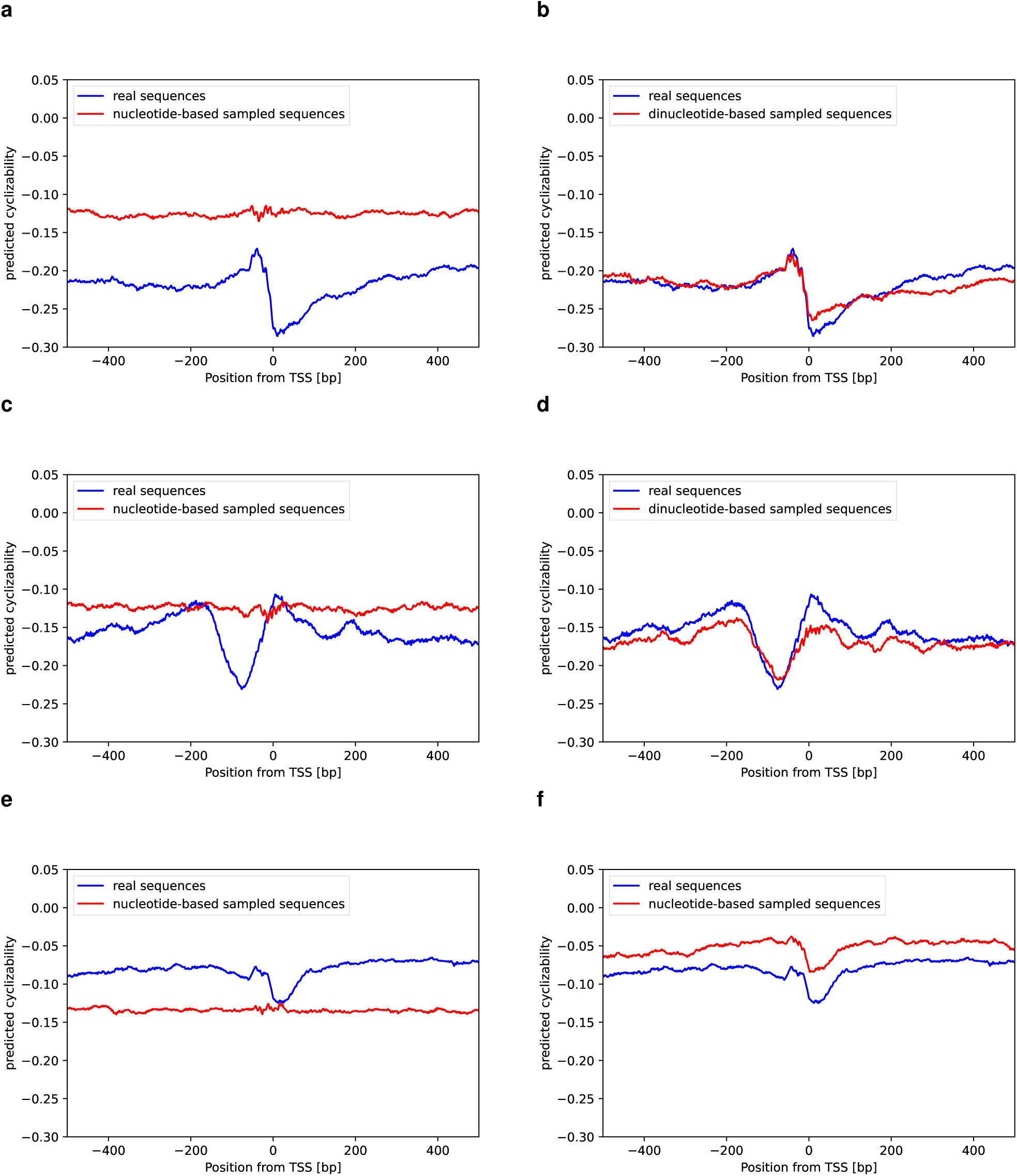
Effects of nucleotide and dinucleotide composition on cyclizability. Mean predicted cyclizability around the TSS of all annotated genes containing a 5’ UTR of compared to sampled sequences from the average base composition at each position and sampled sequences from the average dinucleotide composition at each position in **(a)** and **(b)** *Arabidopsis thaliana*, **(c)** and **(d)** *Saccharomyces cerevisiae*, **(e)** and **(f)** *Homo sapiens*, respectively.

When employing the horizontal shuffling protocol, i.e. shuffling sequence windows of length 50bp horizontally at the level of single bases or dinucleotides), similar observations were made. In the case of mononucleotide random shuffling and using *Arabidopsis thaliana,* as the test species, the characteristic change of cyclizability around TSSs disappeared and the predicted cyclizability was found shifted to increased cyclizability values, i.e. they were predicted more flexible than the actual sequence (Supplementary Figure S1a). By contrast, in the case of dinucleotide shuffling, the signal was barely affected (Supplementary Figure S1b).

Both shuffling-type experiments indicate that dinucleotide composition plays an important role in the intrinsic cyclizability of a given DNA sequence. This agrees with previous studies that linked dinucleotides as key determinants of certain DNA mechanical properties (45, 46) and the study by Khan *et al.* (30), which made similar observations. When comparing the cyclizability signal to the distribution of all 16 dinucleotides at each position normalized to their random expectation, we found that their relative frequencies change markedly around the TSS and that TA frequency, in particular, correlated strongly with the cyclizability pattern found in *Arabidopsis* (Figure 5). In agreement, the correlation between dinucleotide content and measured cyclizability in the random library reported by Basu *et al.* (28) was found strongest for TA (Pearson correlation coefficient, r=0.19). The next two strongest correlated (absolute) dinucleotides, TC (r=-0.09) and GA (-0.081) were found negatively correlated with cyclizability, which also can be observed in the dinucleotide composition around the TSS (Figure 5). Similar observations can be made in *S. cerevisiae* (Supplementary Figure S2) and *Homo sapiens* (Supplementary Figure S3).

**Figure 5.**
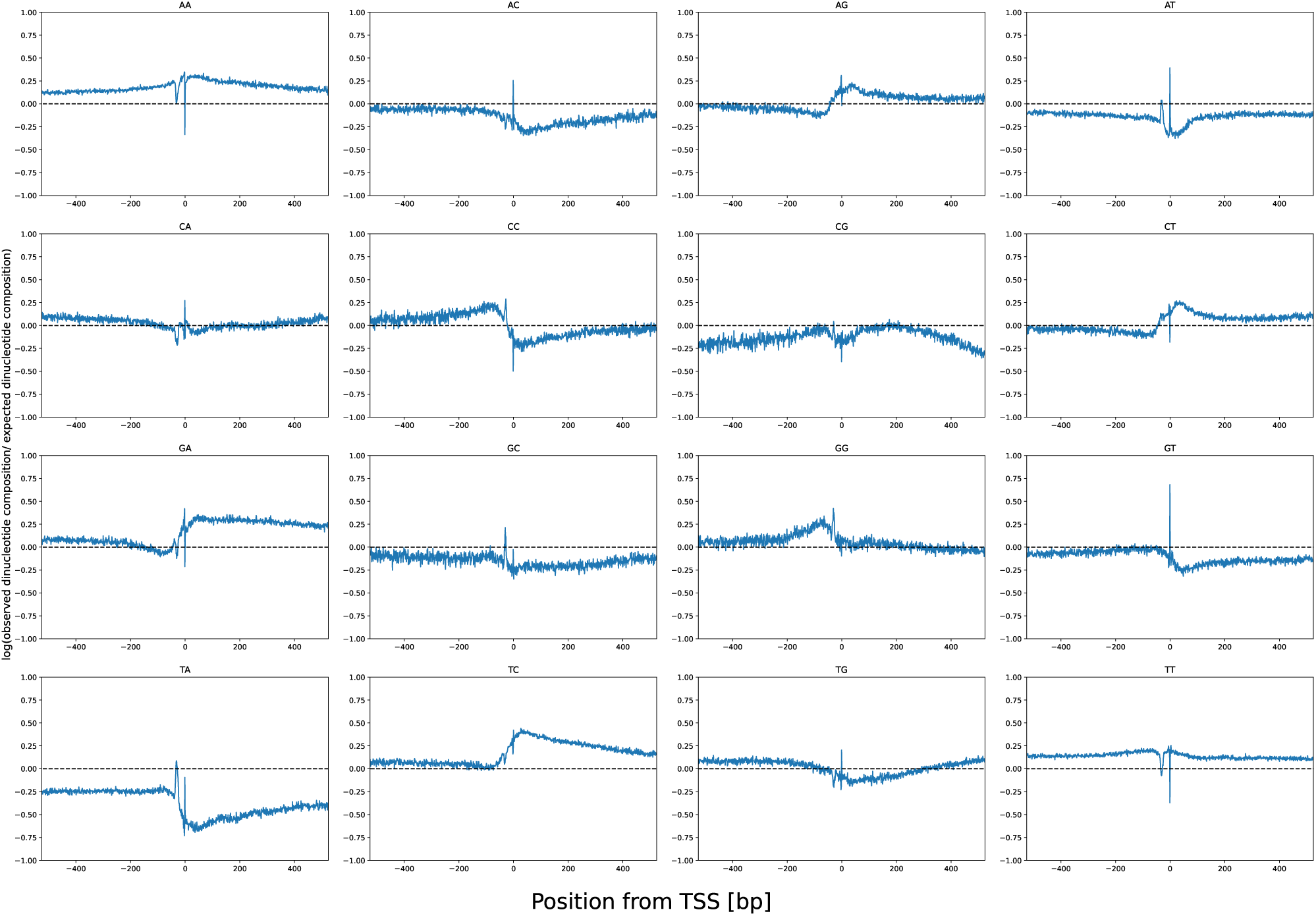
Relative dinucleotide composition around TSSs in *Arabidopsis thaliana*. Natural logarithm of the ratio between the observed dinucleotide composition and the randomly expected dinucleotide composition based on dinucleotide frequency at each position relative to the TSS. Positive/ negative log-ratios imply increased/decreased dinucleotide frequencies relative to random expectation, with the dashed line signifying occurrence as expected.

Two composition peaks, one about 30-40 bp upstream of the TSS and one exactly on the TSS can be observed for most dinucleotides (Figure 5). The latter reflects a high relative occurrence of AT and TG. It seems therefore likely that this peak is largely due to a subset of translation start sites being annotated as TSS. GT, which would be part of the alternate start codon GTG/GUG (47), is also found in high frequencies at that position. The TSS-upstream peak coincides with the reported position of the TATA box motif that is frequently found (∼29% (48)) upstream of TSSs in *Arabidopsis thaliana*.

However, also more complex motifs than dimers are expected to impact DNA mechanics. Using deeplift (38) importance score, we tried to identify potential motifs with TF-modisco (39). However, no motif beyond dimers were identified. Khan *et al.* and using their visible CNN model were able to identify several motifs affecting bendability of a DNA sequence, particularly GAAGAGC and its reverse complement, which was a strong indicator for high flexibility. This effect was confirmed using our own model. Random sequences containing this motif (10000 sequences, compared to shuffled versions of these sequences) were on average significantly more flexible than dinucleotide shuffled versions of the same sequence (Cohen’s d=2.01, p < 0.001).

Taken together, our results indicate that, while dinucleotide composition is a key factor in local DNA flexibility, but higher order motifs may also play an important role.

### Genomic DNA of many eukaryotes is significantly more rigid than expected by random chance

When comparing the rigidity of sequences close to the TSS to shuffled versions (single-base level) of the same sequence in Arabidopsis, we noticed that the original sequences were, on average, more rigid than their randomized versions. Similarly, when we compared predicted cyclizability around the TSS of different species, the investigated vertebrates (mouse and human) showed a consistently higher flexibility (Figure 2i). A similar observation was made by Li et al. (29), who reported that mammals and thermophilic archaea had, in general, a more flexible genome than yeast, *E. coli* and T4 Phage.

We tested whether these observations are statistically significant, and if the genomes of the respective organisms are generally more flexible or rigid than expected by chance. For this, we sampled sequence windows in 50 bp steps from each organism’s genome, respectively, and compared their predicted cyclizability to predicted cyclizability values of the same sequences randomly shuffled (single-base shuffling, five repeats). This revealed that in all eukaryotic organisms investigated, except vertebrates, genomic sequences were significantly more rigid than the randomly shuffled sequences (t-test, p << 0.0001) (Figure 6a). In all non-vertebrate eukaryotes, the effect size between original and randomized sequences ranged from 0.183 in C.elegans to 0.101 in yeast. In the case of vertebrates, the opposite was true, with all of them being, on average, more flexible than randomly shuffled sequences, however with considerably smaller effect sizes, ranging from 0.08 in salmon (*Salmo salar*) to 0.029 in human. When comparing the distributions of differences between genomic vs. shuffled, the largest interspecies difference was found between salmon and C. elegans, with an effect size of 0.3, the smallest between zebrafish and mouse, with an effect size of 0.0026.

**Figure 6.**
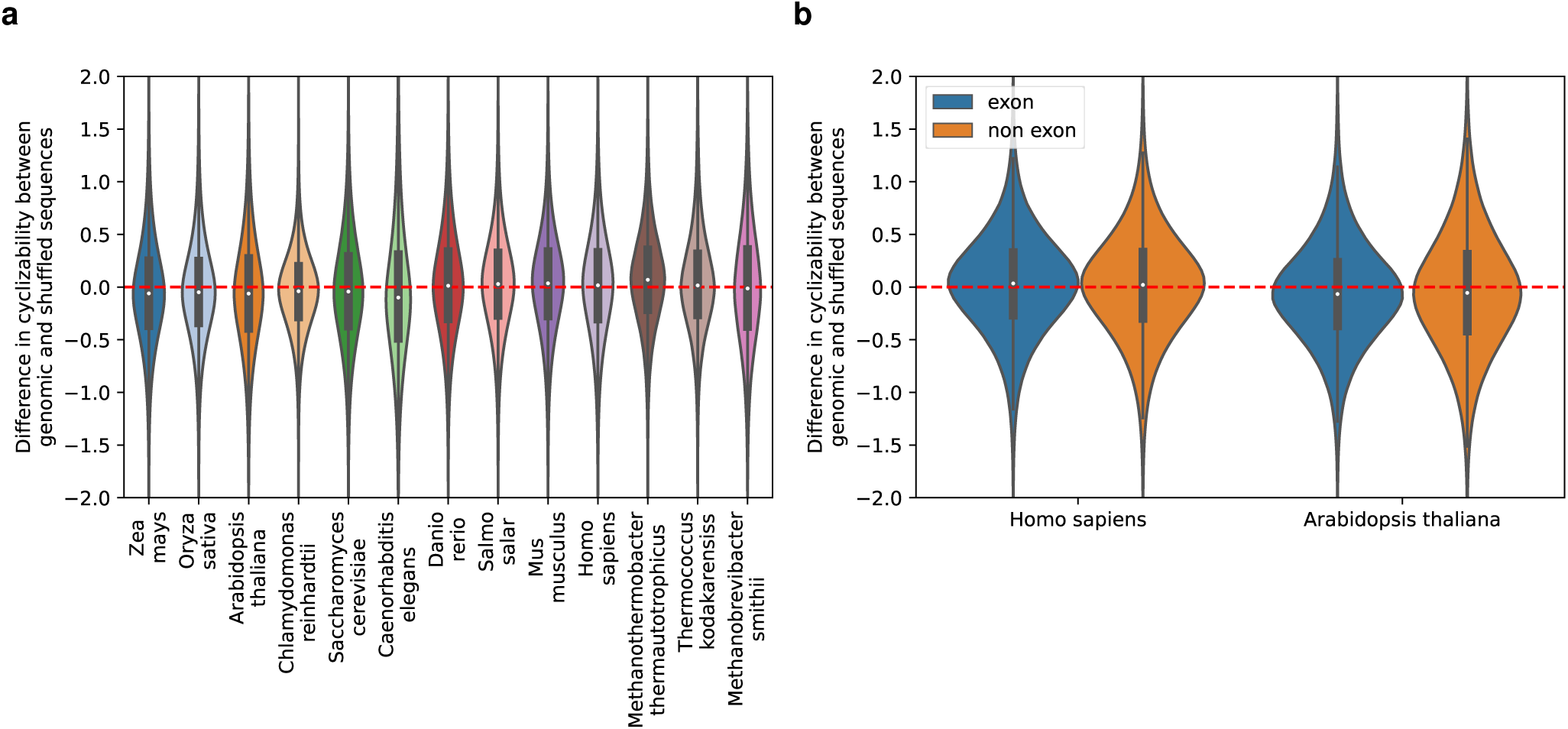
Mean genomic cyclizability of different species relative to randomized versions of the same sequence. **(a)** Distribution of differences between predicted cyclizability of genomic sequences and randomly shuffled versions of the same sequences for different eukaryotes and three archaea (Mth, Tko, and Msm) species. The individual values were derived by predictions of cyclizability of 50bp windows at 50bp step-size (spacing between windows) from the respective genomes, randomly shuffling those sequences five times, and then calculating the difference in cyclizability between shuffled and original sequence. Positive values indicate genomic sequences to be more flexible than their randomized versions, negative values the opposite. **(b)** Distribution of differences between predicted cyclizability of exon/non-exon sequences and randomly shuffled versions of the same sequences for *Homo sapiens* and *Arabidopsis thaliana*. Values were calculated as described under **(a)**.

To exclude composition-based effects, we compared the randomly shuffled sequences to each other. While all differences were significant (p << 0.001), the effect sizes were small, with the largest difference being between *Chlamydomonas reinhardtii* and C. elegans with an effect size of 0.047 and the smallest between maize and rice, with effect size smaller than 0.001. Chlamydomonas was generally an outlier, with the highest mean cyclizability of all shuffled sequences. This is most likely an effect of its very high GC content of Chlamydomonas of around 64% (49) compared to the other species (∼35%-40%). Excluding Chlamydomonas, the highest effect size between randomly shuffled sequences was 0.031. Therefore, it seems likely that the difference in mean cyclizability between genomes is an effect that is unrelated to genome-specific sequence composition.

The occurrence of repetitive DNA-elements could be another possible explanation for differences between species. Vertebrates are known for their large genomes, containing high percentages of repetitive sequences, while most chosen non-vertebrate organisms have relatively short, gene-dense genomes (50). As a test for a potential effect, we compared sequences lying in exons to non-exon sequences in *Arabidopsis* and human, respectively, following the rationale that exons are generally devoid of repetitive sequences. While the respective differences were significant (p < 0.001) due to the large sample size, the very small effect sizes of 0.015 and 0.013 indicated that there were no relevant differences in either species (Figure 6b).

When analyzing three selected archaea species, two thermophiles also investigated by Li *et al.*(29)*, Methanothermobacter thermautotrophicus and Thermococcus kodakarensiss,* and the mammalian commensal *Methanobrevibacter smithii,* we found a strong difference between the species, similar to what we observed in eukaryotes. The two thermophiles (*M. thermautotrophicus* and *T. kodakarensiss*) were significantly more flexible than expected (p << 0.001), with effect sizes of 0.21 and 0.11, respectively. *Methanobrevibacter smithii* however was significantly less flexible than expected (p < 0.05), albeit with a small effect size of 0.013.

### A subgroup of *Arabidopsis thaliana* transcription factors shows distinct DNA rigidity patterns at their binding position compared to non-binding locations of the same motif

The binding of transcription factors (TFs) to DNA is one of the most investigated instances of the role of DNA mechanics. Of particular interest has been the observation that some instances of a TF binding site motif are bound by the respective TF, while for other instances of the same motif, the respective TF shows a much lower affinity. One possible reason could be the local flexibility of DNA, promoting or inhibiting protein binding. Local flexibility is not only dependent on the sequence of the motif, but also the flanking regions. The flanking regions of motifs, while less conserved, play an important role in TF motif recognition and binding (51). Recently, this has been linked to the shape and flexibility of these regions (17, 18).

To investigate this further, we used available DAP-seq data for Arabidopsis TFs (35) and their reported motifs, represented by their position weight matrices (PWM) available on JASPAR, to find bound and unbound motif occurrences in the region 1kbp upstream of any transcription start site (TSS). DAP-seq data were taken as evidence of TF-binding, while absence thereof at respective motif locations were taken as an indication of no binding. For each motif, all bound instances including their -200bp and +200bp flanking region were selected, and a cyclizability profile was predicted. Inspecting the 25 TFs, for which sufficient data was available (Figure 7), we observed that while the majority of TFs showed no clear signal at their binding site with detected binding, others (seven, based on visual inspection: ARID6, ATHB-23, CDF5, NAC017, TCX3, TRB2, and TRp2) have a distinct pattern of predicted rigidity at their binding motif relative to all motif instances. For example, TCX3 shows a distinct pattern of strong negative cyclizability, indicating a preference for rigid binding sites. The opposite of this can be observed when looking at TRP2, which showed a flexible binding site. In this analysis, in order to investigate whether the mechanical properties of the bound regions are actually promoting TF binding, or are only a byproduct of the specific binding sequence, we compared bound motifs to unbound motifs with a q-value less than 0.1 identified by the program FIMO.

**Figure 7.**
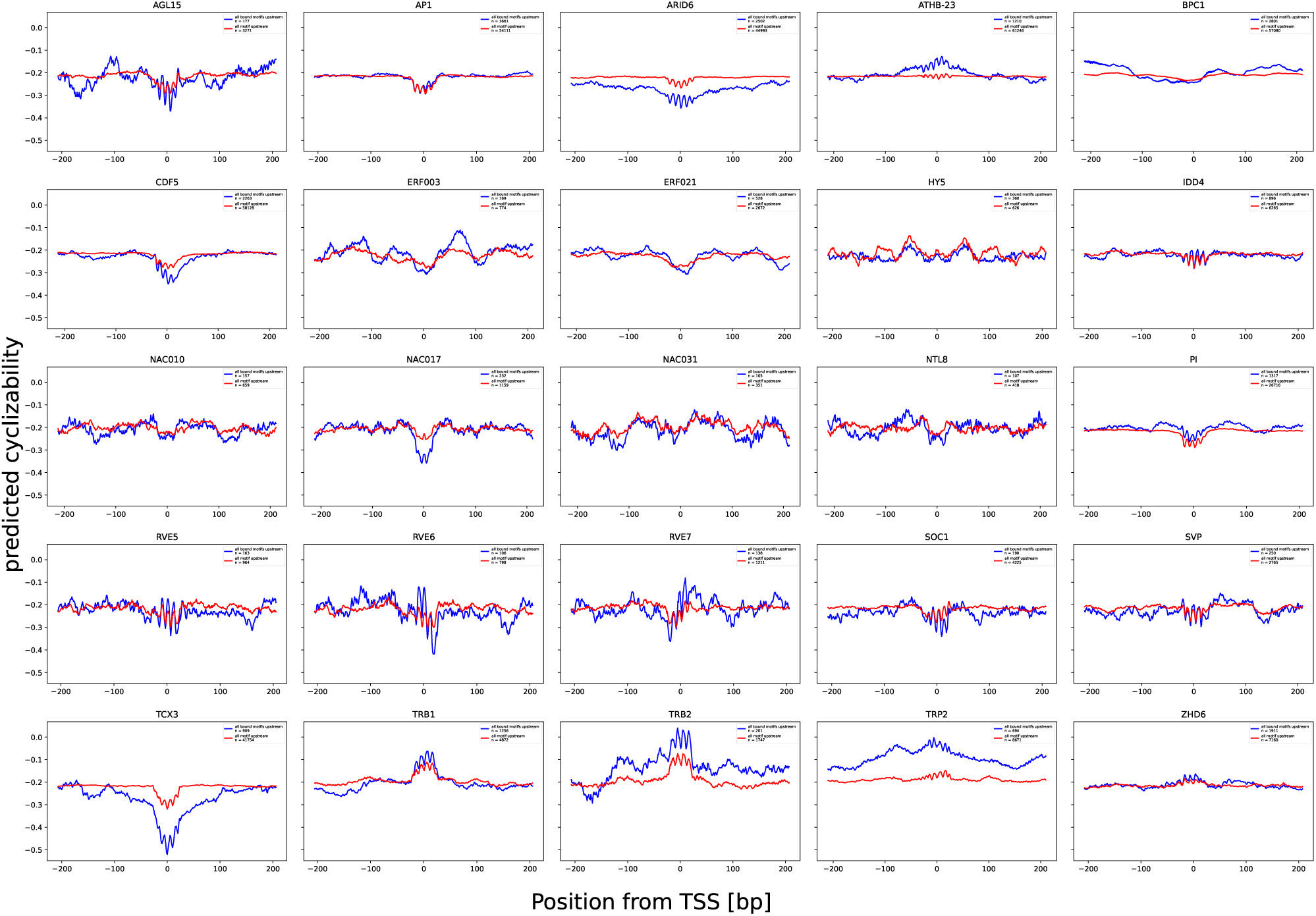
Comparison of bound and unbound instances of transcription factor binding site motifs in gene promoter regions in *Arabidopsis thaliana.* Mean predicted cyclizability of all bound occurrences of transcription factor binding site motifs of a given transcription factor with FIMO q-val score <0.1 (blue profiles) versus all occurrences of the binding motif with the same confidence (red profiles), centered around the binding motif. Only occurrences in 1kbp upstream of annotated TSS were considered in the analysis. TF-binding (bound vs. unbound) was taken as evidenced by DAP-seq support. Presented are all TFs, which met the following criteria: at least 100 found instances of FIMO hits with a q-val below the 0.1 threshold in regions up to 1000bp upstream of any TSS.

Taken together, this indicates that the significance of mechanical properties around the binding site are dependent on the specific TF.

### Methylated positions in *Arabidopsis thaliana* show no preference in local cyclizability

DNA methylation is an epigenetic modification, in which the C5 position of a cytosine is modified by addition of a methyl group. It regulates a wide variety of functions, including gene expression and chromatin regulation. In plants, it can appear in three different contexts, CG, CHG, and CHH. DNA-methylation, in turn, has also been shown to increase DNA flexibility (52). Since Basu et al. were able to show that certain chromatin remodelers were sensitive to local DNA flexibility (28), it is reasonable to assume that DNA methylation might also be influenced by local mechanical DNA properties. Using the methylation calls for *Arabidopsis thaliana* (accession, Col-0) from the 1001 epigenome project (31), we sampled all sequence windows containing an observed methylated position and compared them to random sequence windows with the same context, but no reported methylation. Methylated positions in all three contexts CG, CHG and CHH had a slightly higher cyclizability value than comparable sequences in the same context. However, the effect sizes were small (0.014, 0.005, 0.042, respectively), indicating that these effects are most likely not biologically relevant (Figure 8a). When inspecting all individual base combinations covered under the CHG and CHH motif descriptions, the effects were still small, however, some contexts like CTA and CCT had slightly larger effect sizes of 0.074 and 0.062 (Figure 8b).

**Figure 8.**
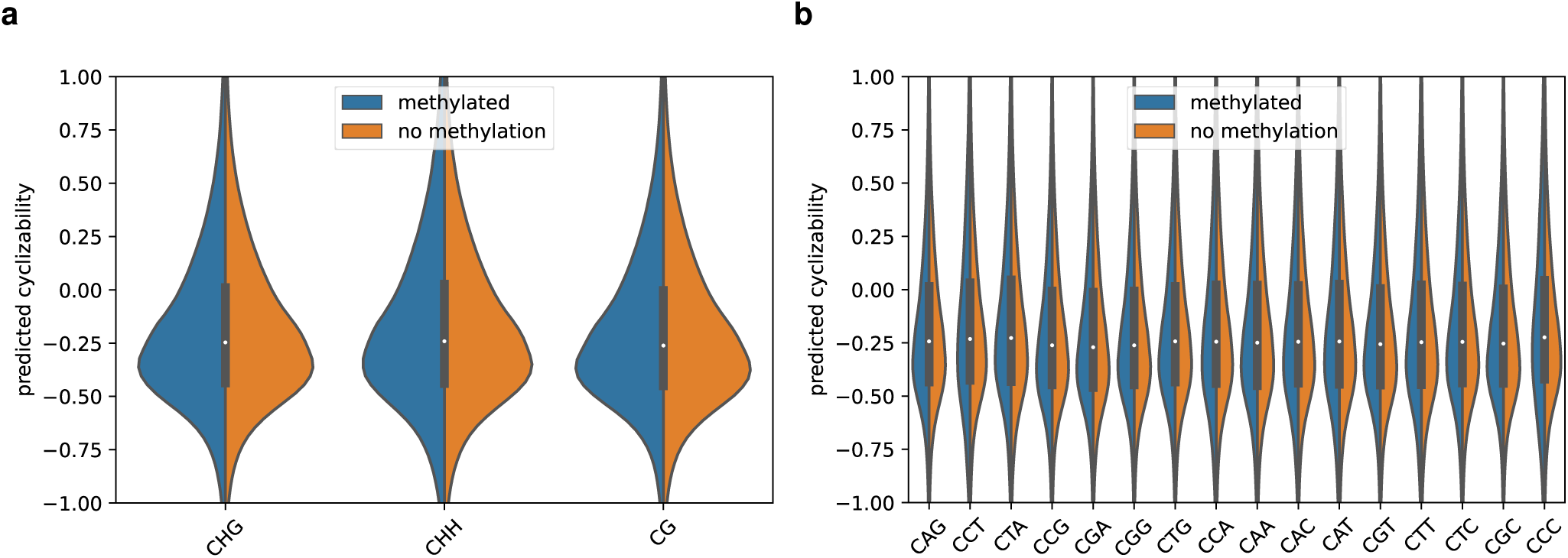
Comparison of cyclizability between methylated and non-methylated positions in *Arabidopsis thaliana.* Comparison of distribution of predicted cyclizability of methylated and unmethylated positions in the same context in the genome of *Arabidopsis thaliana*. Comparison of **(a)** the three main methylation contexts as represented by their consensus motifs: CHG, CHH, and CG, **(b)** all methylation contexts.

### *De novo* mutations and fixed SNPs in *Arabidopsis thaliana* appear more frequently in rigid genomic DNA contexts

A recent paper by Monroe *et al.* revealed that mutations, unlike previously assumed, do not occur at random, but have themselves a bias reflecting natural selection (33). Mutations, accruing *de novo* in the experimental series, were found to occur less often in functional constrained regions of the genome, i.e. gene bodies. This suggests that DNA modifications, DNA accessibility as well as intrinsic properties of certain functional genomic regions could influence the likeliness of *de novo* mutations. They reported that the region upstream of the TSS showed significantly higher mutation rate than the gene body. The mutation rate profile around TSSs behaved very similarly to the predicted cyclizability observed in this study (33).

This led us to surmise that DNA flexibility might be a factor influencing *de novo* mutation rate. Using the provided data from Monroe et al., we sampled all sequence windows containing an observed de novo mutation and compared them to all sequence windows with no reported *de novo* mutation. To exclude any composition and position related effects, we focused gene promoters as a specific genomic region. Considering all sequence windows up to 300bp upstream of a TSS, windows containing *de novo* mutations were significantly more rigid than windows not containing de novo mutations (p < 0.001), with an effect size of 0.146. When comparing all windows with and without *de novo* mutations on Arabidopsis chromosome 1, i.e. without focusing on a specific functional regions, the same trend was observed, with an effect size of 0.134. This implies that rigid regions are generally more susceptible to *de novo* mutations.

Next, we grouped sequence windows into four classes based on presence/absence of particular *de novo* structural variants - no mutation, only single nucleotide polymorphism (SNP) mutation, only indel (insertions/deletions) mutation and SNP and indel mutation. Compared to sequence windows with no mutations, sequence windows of all three mutation classes had lower cyclizability, and are therefore predicted to be less flexible, consistent with the results reported above. However, the strength of the effect varied among the groups (Figure 9a). In the promoter regions, sequence windows with only SNPs had the lowest effect size of 0.108 when compared to windows without mutations. Windows containing only indel mutations had an effect size of 0.167. Lastly, the strongest effect could be observed when comparing windows containing both SNP and indel mutations, with an effect size of 0.309. The results for all windows on chromosome 1 were again comparable, with effect sizes of 0.106, 0.143, and 0.258 for only SNP, only indel and both SNP and indel mutations, respectively.

**Figure 9.**
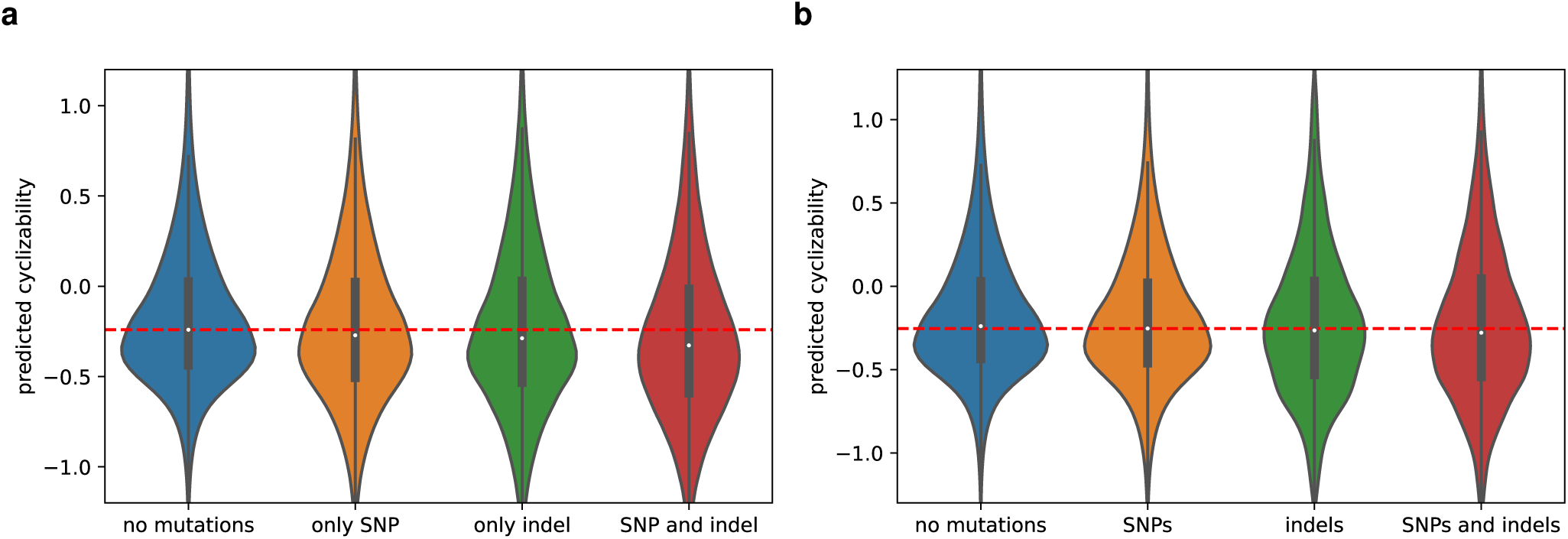
Cyclizability of genomic sequence windows with and without variants in *Arabidopsis thaliana.* **(a)** Comparison of predicted cyclizability between sequences containing no *de novo* mutation, only SNPs, only indel, and both SNP and indel *de nov*o mutations, **(b)** Comparison of predicted cyclizability between sequences containing no fixed mutations (fixed = occurring in naturally occurring accessions, only SNP, only indel, and both SNP and indel mutations according to the 1001 genome project. The dashed lines visualize the genomic median cyclizability value as a reference.

We then investigated whether this effect is observable in fixed SNPs, i.e. naturally occurring SNPs, and short indels, using data from the 1001 genome project (32). When comparing all sequence windows containing mutations to all windows without mutations on chromosome 1 of Arabidopsis, we found that SNP-containing windows were significantly more rigid than non-SNP windows, as observed with *de novo* mutations. The effect size was smaller (0.059) compared to the *de novo* mutation effect sizes, potentially indicating that there is no relevant, or a lesser contribution of mechanical factors in fixed than in *de novo* variants. The different mutation types behave similarly as observed for *de novo* mutations, with effect sizes of 0.058 for SNPs 0.116 for indels and 0.134 for SNPs and indels relative to sequence windows devoid of mutations.

### Sites of high Spo11 oligonucleotide accumulation in *Arabidopsis thaliana* is associated with rigid DNA windows

During meiosis, crossing-over is initiated by double-strand breaks and subsequent DNA repair. These double strand breaks are introduced by Spo11, a relative of topoisomerases (53). Since it has been shown that Spo11-induced breaks are associated with nucleosome depleted regions (54), we suspected that Spo11 might have an affinity for rigid DNA. Using the Spo-11 oligonucleotide mapping in Arabidopsis from Choi et al. (36), we compared those values to the cyclizability values for sequence windows of length 50bp centered around this position. Spo11-oligonucleotides and cyclizability showed no relevant correlation, albeit significant (r=-0.003, p < 0.001). However, due to the high noise of the Spo11 data, we decided to compare the top/bottom two-percentiles of cyclizability sequence windows to the top/bottom two-percentile of Spo11-olignonulceotide score. A Fisher exact test revealed that there is a significantly higher number of sequence windows with high log2(spo11-olignocluetoide/gDNA) and low predicted cyclizability than expected (odds ratio 1.15, p << 0.001). Since TSSs were found associated with low cyclizability (Figure 2), we investigated whether this association was due to an enrichment of high Spo11-oligonucleotide in TSS regions. While the original study by Choi et al. found that Spo11 hotspots were found enriched in the promoter region, we found that the top two-percentile were actually significantly underrepresented in the 1kbp region upstream of the TSS (odds ratio 0.85, p << 0.001).

## Discussion

In this study, we developed, CycPred, a computational artificial neural network prediction model on the basis of available experimental data of genomic sequence cyclizability (28) that proved to perform reliably and fast. Using CycPred we systematically examined how different aspects and characteristic loci of eukaryotic genomes may be affected by DNA flexibility.

Observed for several and diverse species, transcription start sites seem to be associated with pronounced changes of local DNA flexibility (Figure 2). It needs to be asked whether this characteristic profile is itself functionally relevant or a consequence of an overrepresentation of particular sequence motifs signifying TSSs and for reasons other than mechanical properties. In particular, the region immediately upstream of TSSs, the core-promoter, is known to harbor characteristic motifs (such as frequent occurrences of TATA-boxes) and, furthermore, in Arabidopsis was shown to exhibit characteristic dinucleotide orientational preferences (55). Thus, the observed profile of mean cyclizability near TSSs may result for those motifs, and whether the associated flexibility itself is of functional relevance will have to be determined.

We observed that the TSS-cyclizability profile is determined almost entirely by dinucleotide composition (Figure 4, Supplementary Figure S1). Similarly, dinucleotide composition has been found correlated with the specific cyclizability profiles at nucleosome positions (29). Particular dinucleotides have already been reported to impart special mechanical properties. For example, “TA”, frequently referred to as a pyrimidine-purine step, has been implied to be associated with high flexibility (26) and to introduce local kinks (56), possibly involving a switch of base pairing from “Watson-Crick” to “Hoogsteen” (57). Given the TATA-box (literally containing the core motif sequence TATA), the high flexibility observed for regions upstream of the TSS may be explainable. Of note, while on average the profiles obtained for actual and dinucleotide-randomized sequence versions were very similar, their actual mean correlation over pairwise windows was detected at r=0.14 only, i.e. individually, substantial scatter is present.

In the seminal paper on the experimentally determined DNA-cyclizability in yeast (28), which forms the basis of this study, a region of lowered flexibility upstream of TSSs has been reported in yeast, and confirmed here using our computational prediction method (Figure 2e). This region was discussed as a stiff region, in which nucleosome binding/formation is impaired, thereby facilitating TF-binding. Surprisingly, this pattern was not observed in any of the other species examined here (Figure 2). Rather, we observed lowered cyclizability right on, or immediately downstream of the TSS. Assuming the prediction method as well as the TSS annotation to be reliable, this may either point to the yeast genome to be special, or that the stiffness of the TSS itself may be relevant for transcription initiation.

Interestingly, several investigated eukaryotic genomes, but not vertebrates, were significantly more rigid compared to randomized sequences (Figure 6). This effect was largely base-composition-independent. The significance of this observation is not yet clear. It could serve some functional reason, or be the result of some underlying factor unrelated to rigidity, like repetitive sequences. However, there were only minimal differences between exon and non-exon sequences in both Arabidopsis or human (Figure 6b), indicating that repetitive sequences may not be the reason.

While we observed that actual genomic sequences are often more rigid than their randomized versions, with cyclizability taken as a surrogate of flexibility, DNA as a whole still appears to be mechanically very flexible, as was discussed before (23–25). Interestingly, the ∼50bp autocorrelation decay distance of cyclizability (Figure 3) is even shorter than the reported persistence length of DNA, which is around 150bp (58). The cyclization of short DNA sequences (25) can thus not be explained with the standard worm-like chain model from which the persistence length is derived (23). However, to explain many functions of DNA, such as TF binding or other protein DNA interactions, a model allowing higher flexibility of DNA may indeed be biologically very important, and thus, cyclizability real and relevant. Several explanations for the observed discrepancy, like the formation of single-stranded bubbles (59), or kinks (60, 61) have been suggested.

Temperature is a known evolutionary factor shaping genome composition and structure. Thermophilic bacteria and archaea have been reported to have high GC content, which has been linked to thermal adaptation due to higher thermostability and potentially increased DNA repair efficiency (62). Li et al.(29) reported high average cyclizability, and thus high mechanical flexibility, for *Methanothermobacter thermautotrophicus* and *Thermococcus kodakarensiss*, two thermophilic archaea. This can partly be explained with their high GC content of 50% and 52% respectively, since long poly(dA:dT) stretches have been found associated with rigid DNA sections (29, 63, 64). When we compared genomic sequences of those two archaea to randomly shuffled versions of these sequences, we could show that their genome was significantly more flexible than expected by random chance (Figure 6a). However, Chlamydomonas, with an even higher GC content of 64%, has a significantly lower median cyclizability and is more rigid than expected by random chance. Li et al. suggested that the nucleosome structure of thermophile species, wrapping only 60bp around nucleosomes (65), requires their genomes to be more flexible. In contrast to the thermophiles with high flexibility, we found that *Methanobrevibacter smithii*, an archaea commensal, was overall less flexible (Supplementary Figure S4), and showed no strong difference compared with randomized sequences (Figure 6a). Assuming that flexible DNA is required for the tight nucleosome wrapping observed in T. kodakarensiss and M. thermautotrophicus, it is less likely to find this nucleosome organization in M. smithii. This is indicative of the large variability of genome organization found in archaea compared to eukaryotes (66).

Investigation of transcription factor binding showed a distinct preference for rigid/flexible areas of DNA for some transcription factors (Figure 7). By comparing bound and unbound motifs, we could show that some TFs had a more pronounced flexibility motif in bound instances. Since motifs are relatively short sequences (∼6-15bp), the flanking regions have a large impact on the flexibility profile. This agrees with previous reports that the sequence of TF motif flanking regions, while not forming a consistent motif themself, have a large impact on TF binding specificity. Specifically, DNA-shape and flexibility of the flanking regions have been suggested as major contributors to this effect (18–20, 67). It would be interesting to further investigate, which TFs show these preferences, and potentially explore, which mechanism is responsible.

DNA modification by methylation is a key feature of eukaryotic genomes. Our investigation found no strong connection between rigidity and the methylation status of cytosines. This implies that the methyltransferases establishing and maintaining DNA methylation show no particular dependence on the flexibility of DNA, at least in the plant *Arabidopsis thaliana* (Figure 8).

By contrast, *de novo* mutations as reported by *Monroe et al.* (33) have been found located in rigid sequence contexts. Specifically sequence windows containing both SNP and indel mutations were found with increased rigidity (Figure 9). This effect is still observable in reduced capacity in fixed SNPs from the 1001 genome project. This effect could help to explain the different mutation rates observed among different species (68), and also explain why we observed different median cyclizability values among the genomes.

It has been previously reported that DNA bound to nucleosomes accumulate more SNP mutations due to inhibition of certain DNA repair pathways, while linkers showed a higher indel mutation rate. This would be in line with the observation in this study, since linkers have been reported by previous studies to be more rigid (28, 29). Non-homologous end joining (NHEJ) repair of double-strand breaks has been associated with indel mutations (69, 70). It is therefore possible that rigid DNA sections are more susceptible to lesions resulting in double-strand breaks, or more prone to errors during DNA repair. Topoisomerases are enzymes that introduce single-and double-strand breaks into DNA. Their main function is reduction of torsion by reducing positive/negative supercoiling, for example during transcription, which itself could be linked to local DNA mechanics. Therefore, if topoisomerases show a preference for certain local DNA mechanical properties, a higher mutation rate at those positions due to more frequent DNA double strand breaks could be a possible explanation. There have been reports that Topoisomerase 1 is associated with transcription dependent mutagenesis (71, 72). Our investigation of Spo11, a relative of topoisomerase, gave some indication for preference of rigid areas of DNA.

With regard to technical aspects, we developed a computational prediction model, and reported results obtained by applying it, based on experimental data from yeast. For the original data, our prediction accuracy proved very high (r=0.93, with regard to true vs. predicted cyclizability obtained for the reported “random sequence set”, r=0.77 for chromosome V). We then applied the model to other species, i.e. different genomic sequences. In as much as i) DNA is universally the same polymer following the same sequence-structure relationships, ii) the experimental data were obtained for “naked” and synthetic DNA, i.e. without chemical modifications or bound proteins that could be species-specific, and iii) the model was trained on very large sequence datasets, and showed high performance on a random library thus we believe a transfer to other species is plausible and will likely generate valid predictions even if the sequences will differ.

Our results are based on statistical associations, experimental evidence is needed to confirm them as association does not imply causation, and other sequence constraints could partly explain the reported observations.

Furthermore, our results rely on the genomic sequence and annotations to be correct. In particular with regard to the position of transcription start sites, this is, for principle - given alternative start sites - and technical reasons, challenging. However, we believe that the fact that we did observe characteristic profiles around TSSs in all species examined (Figure 2, Figure 5, Supplementary Figures S2, S3), even in view of potential sources of error, suggests that indeed a signal is present.

## Conclusions

Our study explored a range of sequence-structure-function relationships of genomic DNA that may be influenced by local DNA flexibility. We found a characteristic change of flexibility near transcription start sites that was found consistently across multiple species and largely driven by dinucleotide compositional effects. Unlike previously discussed as an indication of facilitated access by transcription factors to gene promoters, no evidence of a generally present region of lowered flexibility upstream of transcription start sites was found. Yet, with regard to transcription factor binding, depending on the actual transcription factor, flanking-sequence-dependent flexibility was observed as a potentially important factor influencing binding. Compared to randomized genomic sequences, depending on species and taxa, actual genomic sequences were globally observed both with increased and lowered flexibility. Lastly, our analysis suggests that crossing-over sites and mutation rate may be linked to local DNA mechanical properties, with rigid areas being more susceptible to SNPs (*de novo* and fixed) and, in particular, indel mutations, and double-strand breaks.

Taken together, our study presents a range of significant correlations between characteristic DNA mechanical properties and genomic features, the significance of which with regard to detailed molecular relevance awaits further experimental and theoretical exploration.

## Availability

CycPred is available at https://github.com/georgback/CycPred.

## Acknowledgements

We wish to thank Ian Henderson for kindly making the Spo11 data available, and Michael Lenhard and Zoran Nikoloski for helpful discussions.

## Authors’ contributions

GB and DW conceived the study, planned the analyses, reviewed and discussed the results, and wrote the manuscript. All computations and code-developments were done by GB.

## Supplementary Figures

**Figure S1.**
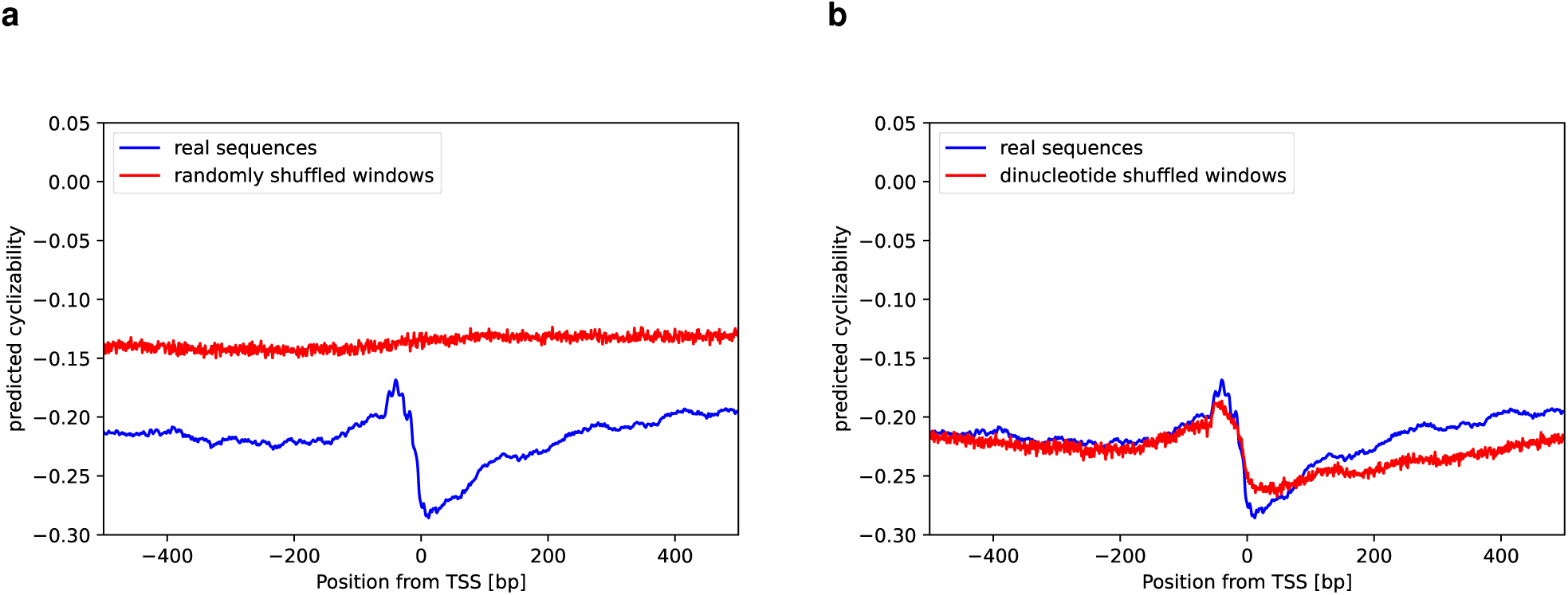
Effects of nucleotide-and dinucleotide-based shuffling on sequences around the TSS. Mean predicted cyclizability around the TSS of all annotated genes containing a 5’ UTR of compared to (a) shuffled versions of all 50 bp windows of the same sequences, (b) dinucleotide shuffled versions of all 50 bp windows of the same sequences in *Arabidopsis thaliana.* The “horizontal” shuffling protocol was applied, i.e. single-bases/ dinucleotides positions were shuffled, unlike drawn from a position-specific probability distribution (“vertical” shuffling).

**Figure S2.**
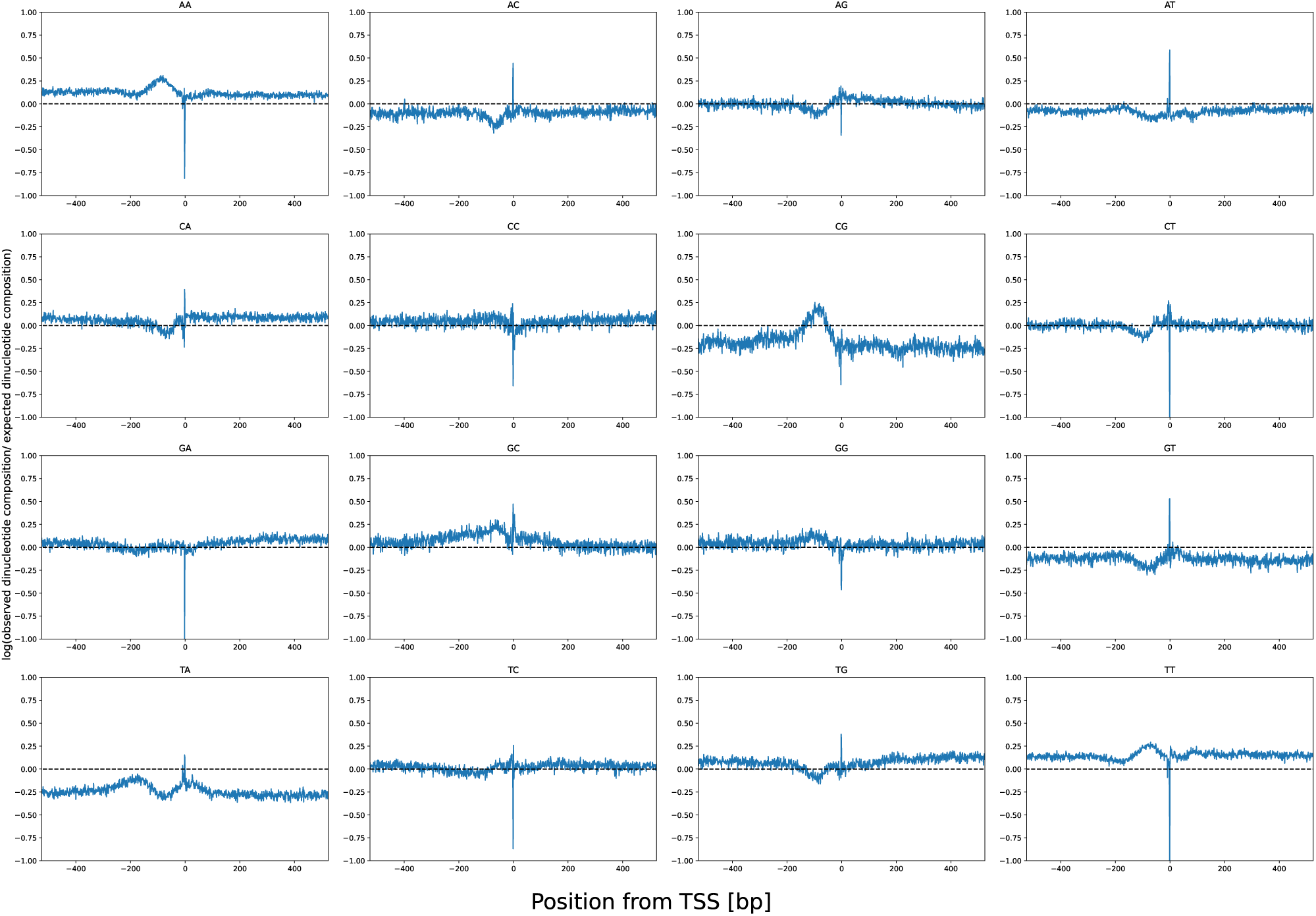
Relative dinucleotide composition around the TSS in *S. cerevisiae.* Natural logarithm of the ratio between the observed dinucleotide composition and the expected dinucleotide composition based on nucleotide frequency at each position in *Saccharomyces cerevisiae*.

**Figure S3.**
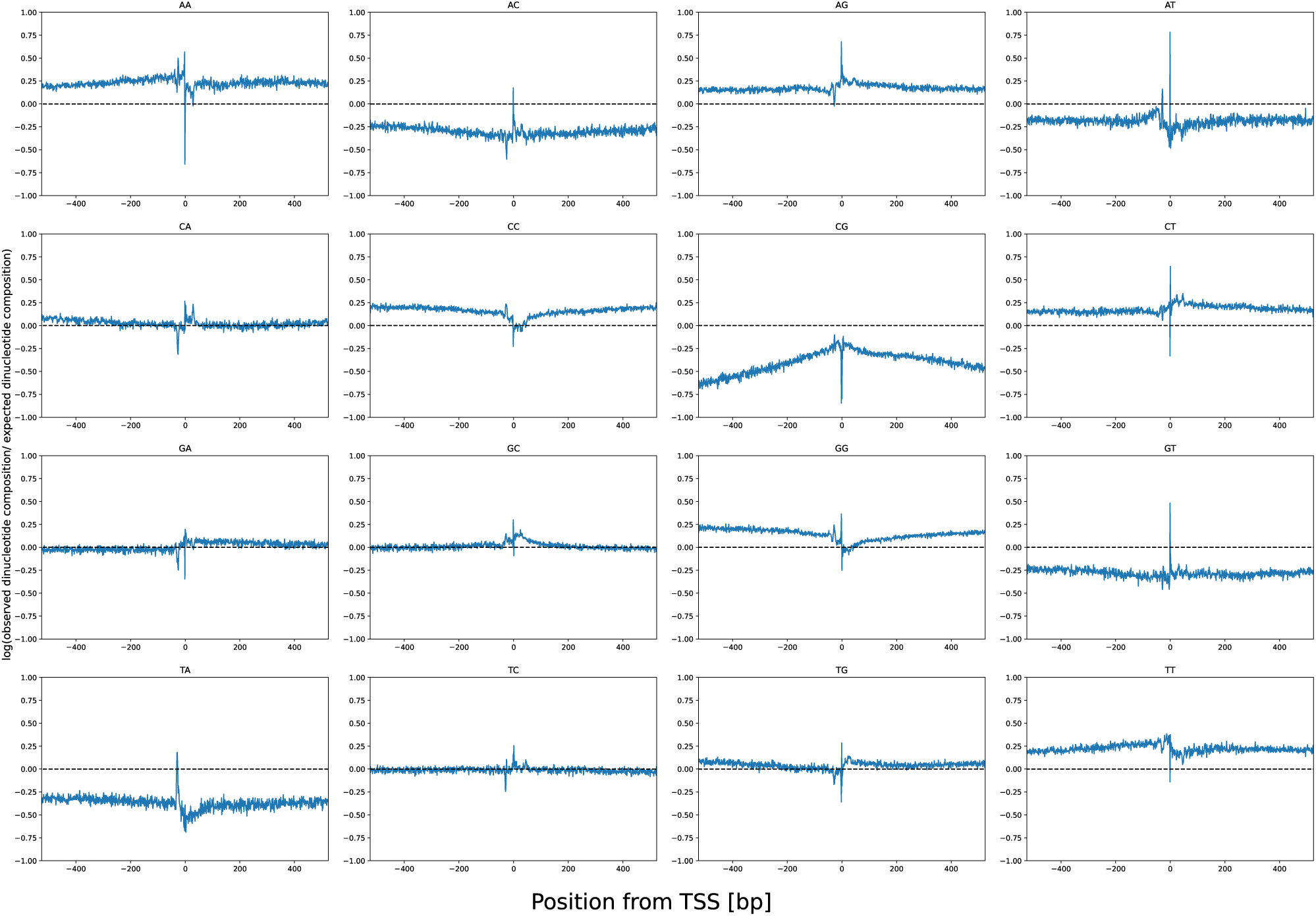
Relative dinucleotide composition around the TSS in *Homo sapiens.* Natural logarithm of the ratio between the observed dinucleotide composition and the expected dinucleotide composition based on nucleotide frequency at each position in *Homo sapiens*.

**Figure S4.**
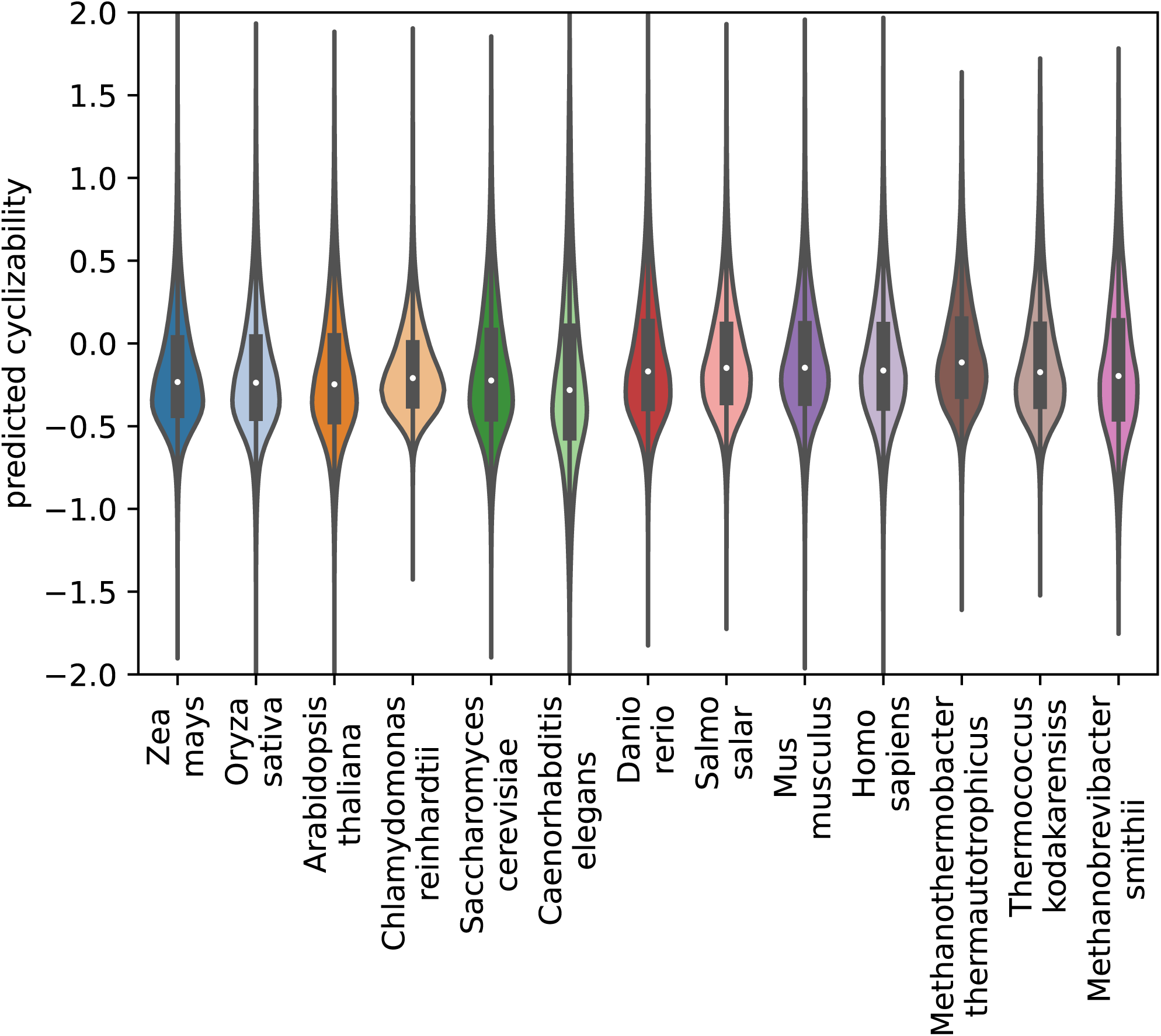
Distribution of predicted cyclizability among different species.

