## Supplementary for "Predictions of DNA mechanical properties at a genomic scale reveal potentially new functional roles of DNA-flexibility"

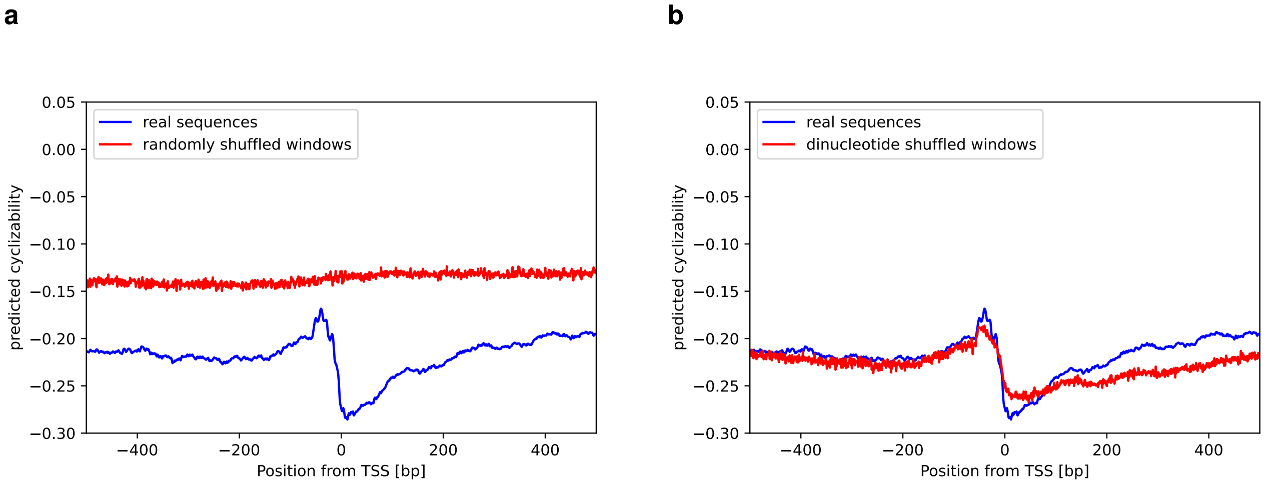


**Figure S1. Effects of nucleotide- and dinucleotide-based shuffling on sequences around the TSS**
Mean predicted cyclizability around the TSS of all annotated genes containing a 5’ UTR of compared to **(a)** shuffled versions of all 50 bp windows of the same sequences, **(b)** dinucleotide shuffled versions of all 50 bp windows of the same sequences in *Arabidopsis thaliana*. The “horizontal” shuffling protocol was applied, i.e. single-bases/ dinucleotides positions were shuffled, unlike drawn from a position-specific probability distribution (“vertical” shuffling).

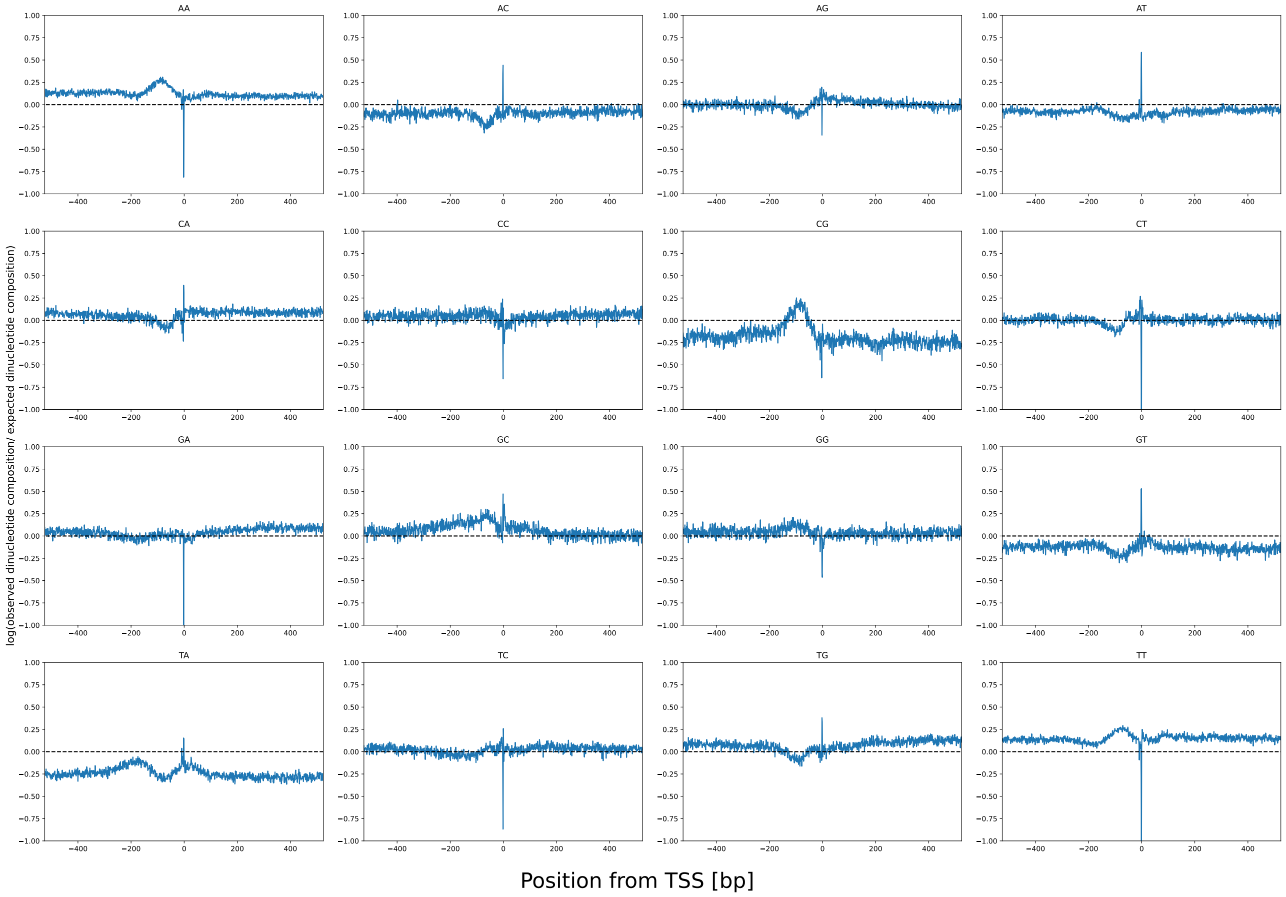


**Figure S2. Relative dinucleotide composition around the TSS in *S. cerevesia*.**
Natural logarithm of the ratio between the observed dinucleotide composition and the expected dinucleotide composition based on nucleotide frequency at each position in *Saccharomyces cerevisiae*

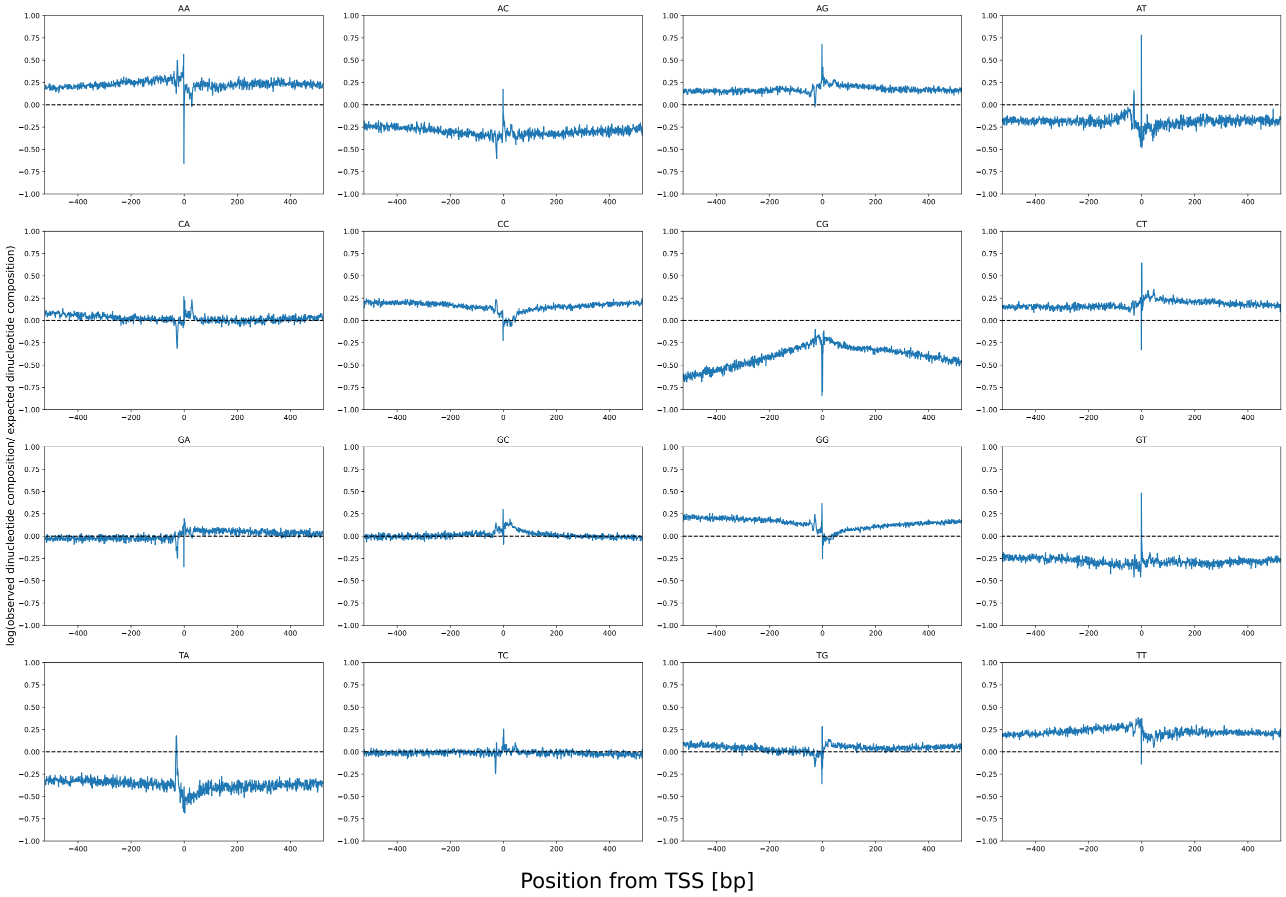


**Figure S3. Relative dinucleotide composition around the TSS in *Homo sapiens*.**
Natural logarithm of the ratio between the observed dinucleotide composition and the expected dinucleotide composition based on nucleotide frequency at each position in *Homo sapiens*

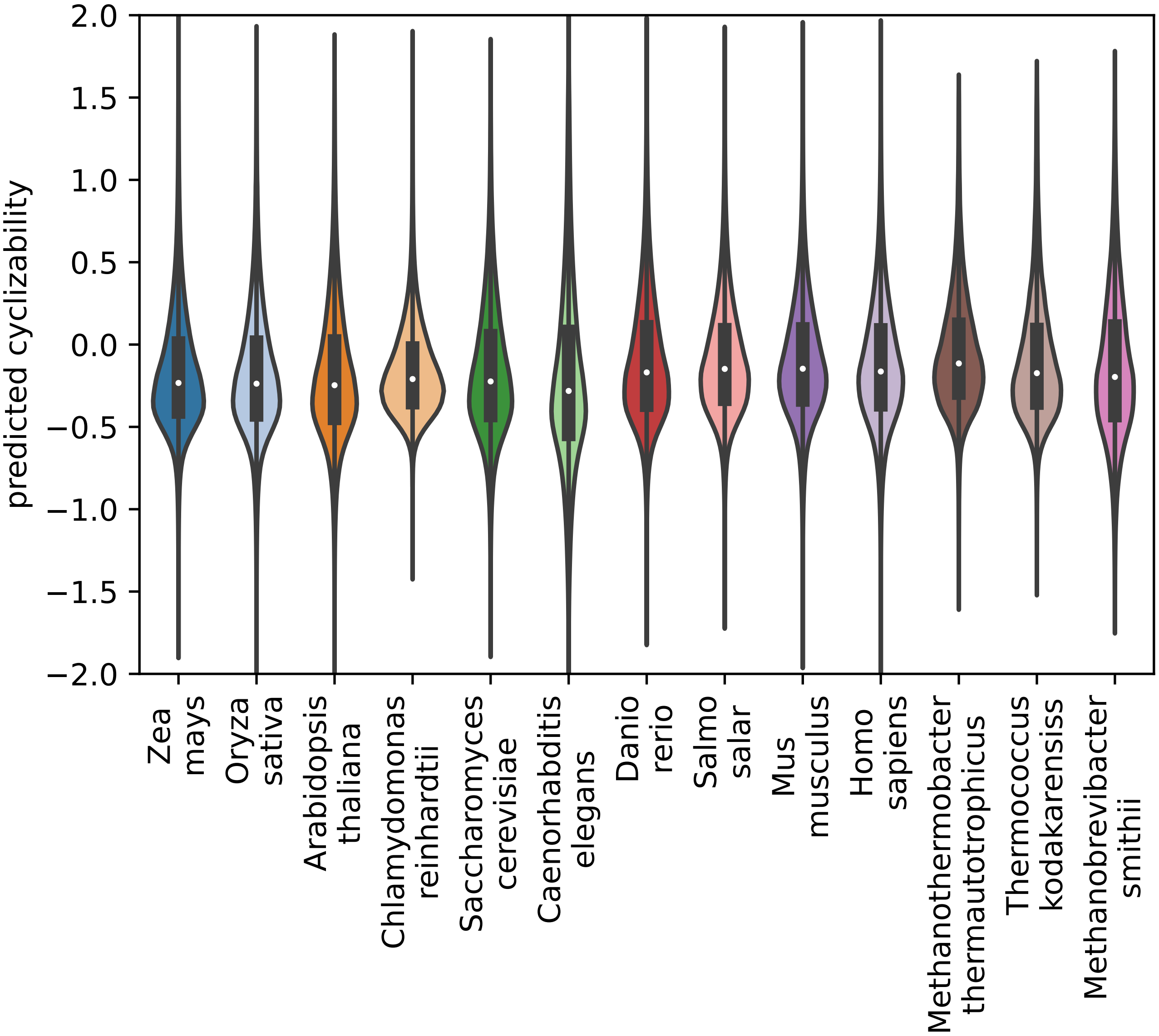


**Figure S4. Distribution of predicted cyclizability among different species**
